# Ddp1 cooperates with Ppx1 to counter a stress response initiated by non-vacuolar polyphosphate

**DOI:** 10.1101/2022.02.02.478728

**Authors:** Liam McCarthy, Iryna Abramchuk, Gamal Wafy, Alix Denoncourt, Mathieu Lavallée-Adam, Michael Downey

## Abstract

In diverse cells from bacterial to mammalian species, inorganic phosphate is stored in long chains called polyphosphates (polyP). These near universal polymers, ranging from 3 to thousands of phosphate moieties in length, are associated with molecular functions including energy homeostasis, protein folding, and cell signaling. In many cell types, polyphosphate is concentrated in subcellular compartments or organelles. In the budding yeast *S. cerevisiae,* polyP synthesis by the membrane-bound VTC complex is coupled to its translocation into the lumen of the vacuole, a lysosome-related organelle, where it is stored at high concentrations. In contrast, ectopic expression of bacterial polyphosphate kinase, PPK, results in the toxic accumulation of polyP outside of the vacuole. In this study, we used label-free mass spectrometry to investigate the mechanisms underlying this toxicity. We find that PPK expression results in the activation of a stress response mediated in part by the Hog1 and Yak1 kinases, and Msn2/Msn4 transcription factors. This response is countered by the combined action of the Ddp1 and Ppx1 polyphosphatases that function together to counter polyP accumulation and downstream toxicity. In contrast, ectopic expression of previously proposed mammalian polyphosphatases did not impact PPK-mediated toxicity in this model, suggesting either that these enzymes do not function directly as polyphosphatases *in vivo* or that they require co-factors unique to higher eukaryotes. Our work provides a mechanistic explanation for why polyP accumulation outside of lysosome-related organelles is toxic. Further, it serves as a resource for exploring how polyP may impact conserved biological processes at a molecular level.

## INTRODUCTION

Inorganic phosphates can be joined in linear chains that range from 3 to 1000s of residues in length. These polymers, called polyphosphate or ‘polyP’, have been implicated in diverse biological processes spanning both prokaryotic and eukaryotic species. In the budding yeast *S. cerevisiae,* polyP is synthesized by the <u>v</u>acuolar transporter <u>c</u>haperone (VTC) complex^[1]^. The VTC complex is composed of the catalytic subunit Vtc4, along with Vtc1 and one of either Vtc2 or Vtc3^[1]^. Cells mutated for Vtc4, Vtc1, or both Vtc2 and Vtc3 have no detectable polyphosphate^[2]^. An accessory subunit called Vtc5 increases the activity of the complex but is not strictly required for polyP synthesis^[3]^. The majority of the VTC complex localizes to the membrane of the yeast vacuole, a type of lysosome-related organelle (LRO)^[1]^. The VTC complex spans the vacuolar membrane and synthesises polyP chains by hydrolyzing the gamma phosphate of cytoplasmic ATP, adding it to growing polyP chains that are simultaneously translocated into the vacuole lumen^[4]^. Vacuolar polyP comprises the majority of total cellular polyP stores exceeding 10-20 % of the dry weight^[2, 5]^. Other areas of the cell beyond the vacuole, including the plasma membrane, cytoplasm, and mitochondria have lower levels of polyP. PolyP is also found in the nucleus, although the reported levels vary drastically^[2]^. These variations are likely due to differences in growth conditions, as well as differences in the method of sample preparation and detection. Whether polyP outside of the vacuole is synthesized locally or somehow transported from polyP stores that first accumulate in the vacuole lumen is unknown. Although some VTC proteins localize in part to the plasma membrane and/or the endoplasmic reticulum^[1, 6]^, it is unknown if these pools of VTC proteins participate in polyP synthesis.

The subcellular enrichment of polyP is conserved in higher eukaryotes^[7]^. For example, in human myeloma cells^[8]^, and various cancer cell lines treated with cisplatin^[9]^, polyP accumulates in the nucleolus. In platelets it is concentrated in LROs called dense granules^[10]^. The localization of polyP to such hotspots likely promotes specific functions. In yeast, vacuolar polyP is important for phosphate homeostasis^[11]^ and vacuole fusion^[12]^. PolyP released from dense granules following injury interacts directly with specific clotting factors to promote blood coagulation^[10, 13]^. In contrast, less is known about whether there are physiological reasons why polyP in some areas of the cell must be kept low.

Unlike the VTC complex, ectopic expression of *Escherichia coli* (*E. coli*) polyphosphate kinase (called PPK) in yeast results in accumulation of polyP outside of the vacuole and this correlates with slow growth, altered cellular morphology, and dramatic sensitivity to the translation inhibitor cycloheximide and mTOR inhibitor rapamycin^[4, 14]^. The amount of polyP made by PPK is at most 20 % of that produced by VTC^[4]^, and PPK-induced toxicity is observed both in wild-type and *vtc4*Δ cells^[14]^. Together, these data point to the location of PPK- synthesized polyP, rather than changes in the total amount, as the driving force behind the observed toxicity. Finding the mechanisms underlying this toxicity will provide a foundation for better understanding the spatial and temporal regulation of polyP metabolism in eukaryotic cells.

Here, we use label-free mass spectrometry to determine why PPK expression and the concomitant accumulation of non-vacuolar polyP is toxic in yeast. Our results demonstrate that PPK-expressing cells remodel their proteome with changes to protein networks controlling fundamental aspects of metabolism. In follow-up work, we present evidence that a Yak1/Hog1-Msn2/4 pathway limits viability in these cells. Finally, we demonstrate that polyP accumulation outside the vacuole is countered by the combined activities of the endopolyphosphatase Ddp1 and the exopolyphosphatase Ppx1 that work together to degrade long-chain polyP synthesized by PPK. Our results provide an explanation for why polyP is sequestered, elaborate functions for polyphosphatases outside the vacuole, and suggest new avenues for exploring polyP functions at the molecular level.

## RESULTS

### Remodelling of the yeast proteome in response to ectopic PPK expression

To determine why PPK expression and the accumulation of non-vacuolar polyP is toxic in yeast, we used label-free mass spectrometry to survey changes to the proteome in strains expressing PPK versus empty vector control. Our initial experiments were carried out in a *vtc4*Δ background, which enabled us to confirm the production of the polyP in each isolate used for analysis, as judged by the lysine polyphosphorylation of Fpr3 protein^[14]^ (**Supplementary Fig. 1**). The polyP made by PPK in yeast was of the long-chain type, similar to that produced by *E. coli* under starvation (**Fig. 1A**). Total protein from 5 isolates of control and 5 isolates of PPK-expressing cells was isolated using an SDS-lysis protocol prior to trypsinization and MS/MS analysis using a Thermo Scientific Q Exactive Plus Orbitrap mass spectrometer (**Fig. 1B**). As expected, peptides from bacterial PPK were detected only in the PPK-expressing strains (**Supplementary Table S1**). At an FDR-adjusted *p*-value (*p*) < 0.05 level, 8 proteins were deemed significantly upregulated, while 6 were downregulated (**Fig. 1C and 1D**). An additional 24 proteins showed up- or downregulation with an associated statistical significance of 0.05 ≤ *p* ≤ 0.1 (**Fig. 1C**). Finally, there were 2 proteins that were detected in all five replicates of the empty vector control, but not in any replicates of the PPK-expressing strains (**Fig. 1D and Supplementary Table S1**).

**Figure 1.**
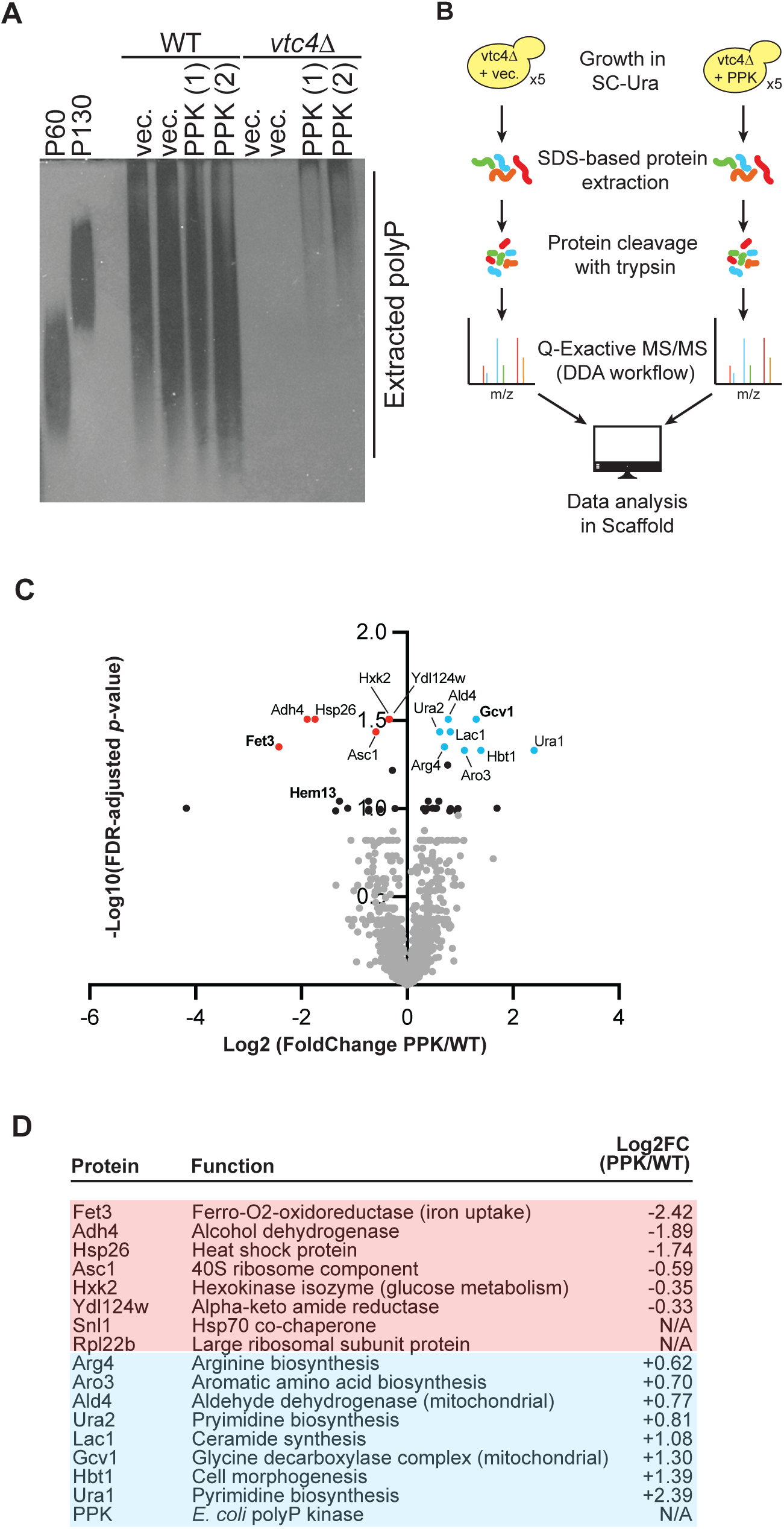
The proteome of PPK-expressing yeast. **A)** PPK expression in yeast allows for production of polyP outside of the vacuole. Polyphosphate was extracted from the strains expressing the indicated constructs and visualized using DAPI staining following separation on 15.8 % acrylamide TBE-Urea gel. **B)** Workflow for data dependent-based label-free mass spectrometry analysis. **C)** Volcano plot with proteins up- or downregulated following PPK expression in a *vtc4*Δ background. Red proteins are downregulated (adjusted *p*-value < 0.05), blue proteins are upregulated (adjusted *p*-value < 0.05). Black proteins are those with adjusted *p*-value ≤ 0.1. **D**) Table of top mis-regulated proteins identified with an adjusted *p*-value < 0.05. Images representative of n ≥ 3 experiments. **Also see Supplementary Figure 1.**

### Vacuolar polyP does not impact the cellular response to PPK expression

To validate our proteomics dataset, we expressed up- or downregulated proteins as C-terminal 3FLAG fusions from their endogenous loci and transformed these strains with control or PPK-expressing vectors. We first carried out this analysis in a *vtc4*Δ background, mirroring the conditions used for our mass spectrometry experiments. For differentially expressed proteins with a *p* ≤ 0.1, we confirmed 6 of 6 upregulations chosen for analysis (e.g. Gcv1) and 3 of 4 downregulations chosen for analysis (e.g. Fet3, Hem13), for an overall confirmation rate of 90 % (**Fig. 2A and Supplementary Table S2**). The validation of protein changes in both the *p* < 0.05 and *p* ≤ 0.1 ranges supports our use of the *p* ≤ 0.1 cut-off for downstream bioinformatics analyses (see below). Notably, we also confirmed 5 of 6 differentially expressed proteins beyond the *p* = 0.1 value range (**Supplementary Table S2**), highlighting the overall high quality of our dataset and the stringency of our statistical approach.

**Figure 2.**
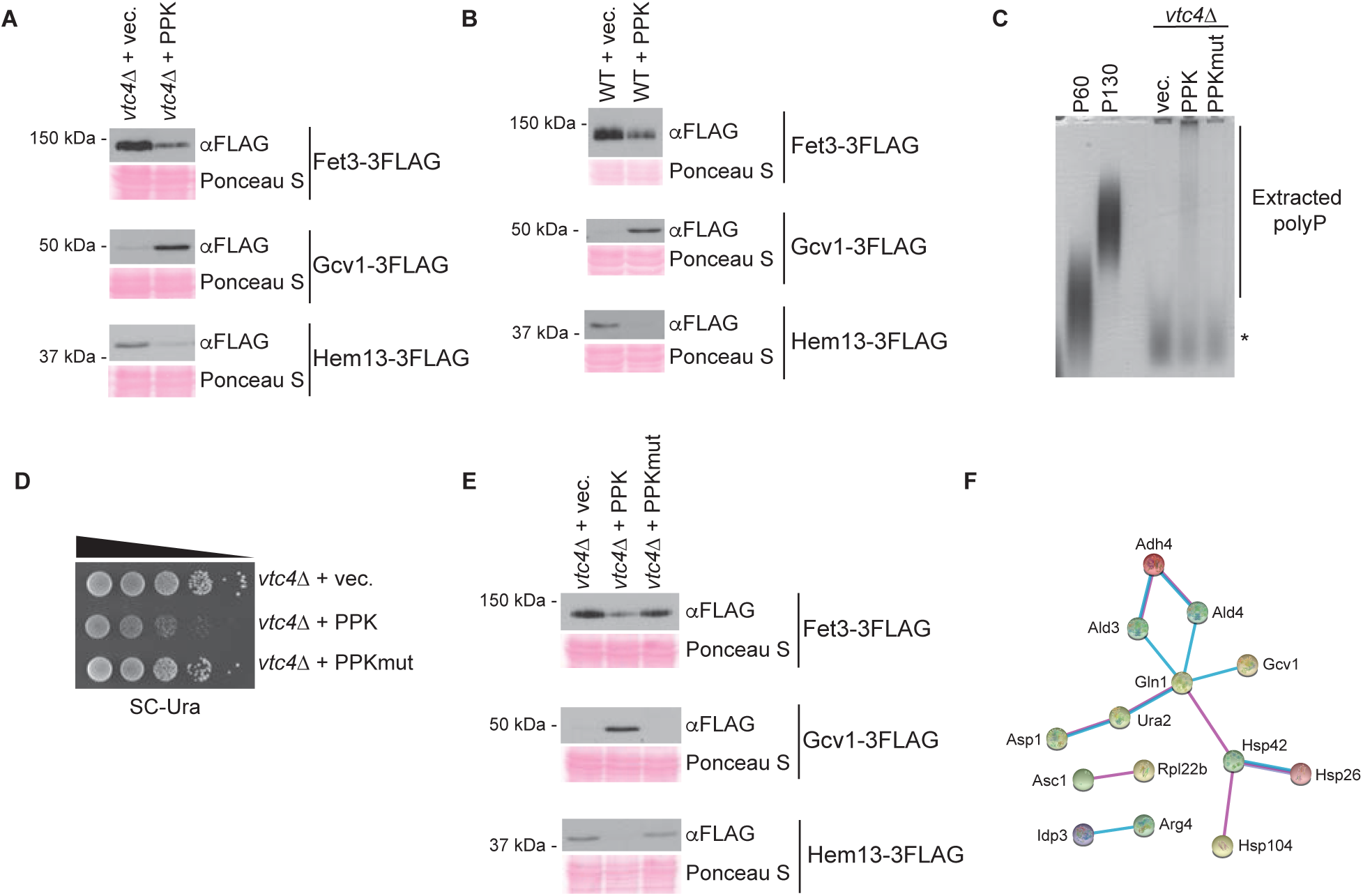
The response to PPK expression is not impacted by the presence of vacuolar polyP. **A)–B)** Protein extracts from the indicated strains were prepared using a TCA-lysis protocol and separated on SDS-PAGE gels. Separated proteins were transferred to PVDF membrane prior to probing with anti-FLAG antibody to detect candidate proteins. Ponceau S stain of the PVDF membrane is shown as a loading control. **C)** Polyphosphate was extracted from the strains expressing the indicated constructs and visualized using toluidine blue staining following separation on a 15.8 % acrylamide TBE-Urea gel. The asterisk indicates non-specific staining of an unknown product that is not polyP. **D)** 10-fold dilutions of the indicated strains were spotted on SC-Ura and incubated for 48 hours at 30 °C. **E)** Protein extracts from the indicated strains were analyzed as described in (A). **F)** STRING analysis locates mis-regulated proteins in sub-networks of physically interacting proteins. Line colours: magenta: experimental evidence, cyan: database evidence, purple: database evidence (specifically protein homology). Images representative of n ≥ 3 experiments. **Also see Supplementary Figure 2.**

To test if these changes are influenced by the presence of vacuolar polyP, we repeated our western-blotting analyses in a wild-type strain (i.e. *VTC4*+), with a confirmation rate of 86 % (*p* ≤ 0.1, **Fig. 2B and Supplementary Table S2**). This observation is consistent with our previous work demonstrating that PPK expression sensitizes cells to rapamycin and cycloheximide, regardless of *VTC4* status^[14]^. As suggested^[4]^, PPK is not simply replacing polyP that is missing in *vtc4*Δ cells. Instead, non-vacuolar polyP appears to have unique consequences for the cell. Finally, to test if the catalytic activity of PPK was required for the observed effects, we generated a mutant version of the enzyme where three histidine residues shown or proposed to be required for catalytic activity^[15–17]^ were mutated to alanine. This mutant enzyme did not produce detectable polyP *in vivo (***Fig. 2C**). Further, it did not result in the same slow growth defects seen for wild-type PPK expression (**Fig. 2D**). For 8 of 8 candidates tested (*p* ≤ 0.1), mutant PPK did not impact protein levels (**Fig. 2E and Supplementary Table S2**). Altogether, our data confirm that PPK-expression impacts polyP levels independently of vacuolar polyP. Finally, we note that proteins encoded by the phosphate (PHO) regulon (e.g. Pho8, Pho86, Vtc2)^[18]^ did not show significant changes in PPK-expressing cells (**Supplementary Table S1**). Therefore, the effects of PPK expression are not easily explained by changes in phosphate signaling.

### Mis-regulation of proteins involved in fundamental metabolism in PPK-expressing cells

To identify molecular functions impacted by PPK expression we carried out a Gene Ontology^[19]^ enrichment analysis of mis-regulated proteins (*p* ≤ 0.1). We found a statistically significant enrichment (FDR-adjusted *p*-value = 0.036) of GO:0006082 – organic acid metabolic process (**Supplementary Table S1**). Mis-regulated proteins belonging to this group include those involved in glucose metabolism (Hxk2, Tdh3), B vitamin synthesis (Snz1, Fol2), and amino acid biogenesis or turnover (e.g. Aro3, Gcv1, Gln1)^[20]^. These changes suggest that polyP accumulation outside of the vacuole influences important aspects of primary metabolism. Using STRING^[21]^ (Search Tool for the Retrieval of Interacting Genes/Proteins) to model protein interactions, we found that mis-regulated proteins involved in amino acid metabolism shared physical interactions with each other, and with 3 heat shock proteins (Hsp26, Hsp42, Hsp104)^[20]^ that were also significantly mis-regulated with PPK expression (**Figure 2F**). The identification of heat shock proteins in this network is interesting in light of the proposed role of polyP itself as a molecular chaperone (See Discussion)^[22]^. Other heat shock or ‘quality control’ proteins mis-regulated in our dataset include Hsp12, Hsp104, and Snl1, a co-chaperone for Hsp70^[20]^.

PolyP can function as an ion chelator, and recent work from the Jakob lab suggests that it can act as an inhibitor of the Fenton reaction, in which Fe^2+^ reacts with hydrogen peroxide to form reactive hydroxyl radicals^[23]^. Interestingly, we noted that a number of proteins mis-regulated with PPK expression are either involved in ion transport (e.g. Fet3, which oxidizes Fe^2+^ to Fe^3+[24]^) or require divalent ions as co-factors (e.g. Adh4, which requires zinc for its catalytic activity^[25]^). Genes encoding other mis-regulated proteins confer resistance to metal ions (e.g. *HIS1*^[26]^), or are induced in response to metal deficiency (e.g. *ADH4*, induced in response to low zinc^[27]^). While the toxicity associated with PPK expression was not impacted by the addition of various divalent cations (Fe^2+^, Mg^2+^, Zn^2+^) to growth media (**Supplementary Fig. 2**), we cannot yet rule out that disruption of ion homeostasis impacts proteome changes independently of the overt toxicity observed in PPK-expressing cells.

### An Msn2/Msn4–dependent signaling pathway limits viability in PPK–expressing cells

We next used the Yeast Search for Transcriptional Regulators and Consensus Tracking (YEASTRACT^[28]^) tool to identify candidate transcription factors involved in the regulation of genes encoding proteins up- or down-regulated in our dataset (**Fig. 3A**). We were particularly interested in Msn2 (regulating 85 % of hits) and its paralog Msn4 (regulating 74 % of hits) for two reasons. First, these transcription factors are involved in the response to diverse cellular stresses via the activation of stress response element genes^[29, 30]^ (**Fig. 3B**). Second, the constitutive activation of Msn2 is associated with slow growth^[31, 32]^. We reasoned that mis-regulation of Msn2 and/or Msn4 impact cell growth and/or division in PPK-expressing cells. Indeed, deletion of *MSN2* and *MSN4* in PPK-expressing strains improved growth in serial dilution assays (**Fig. 3C**), suggesting that these transcription factors hinder growth in the presence of non-vacuolar polyP. We next examined changes to Msn2 via western blotting. For these experiments, we developed an integrated GAL-inducible version of the PPK expression system (**Fig. 3D**). This allowed for better control over PPK expression and prevented strains from adapting to the presence of non-vacuolar polyP. The PPK construct contains an N-terminal HA epitope tag, which allowed us to monitor expression in mutant strains. Notably, GAL-induced HA-PPK expression resulted in growth defects in both *vtc4*Δ and WT strain backgrounds (**Fig. 3D**). We further validated our GAL-inducible system by demonstrating that Fet3-3FLAG is downregulated and Gcv1-3FLAG is upregulated following HA-PPK expression (**Fig. 3E**), mirroring what was observed for the plasmid-based expression system used in previous experiments. Loss of PKA dependent Msn2-NLS phosphorylation is associated with its nuclear localization and transcriptional activation of stress response element genes (**Fig 3B**)^[33]^. HA-PPK-expressing cells had lower levels of this phosphorylated form of Msn2, which can be fortuitously detected using an antibody against human pCREB^[33, 34]^ (**Fig. 3F**). Notably, total levels of total Msn2 were also reduced with PPK expression (**Fig. 3F**), an observation that may seem at odds with the idea of enhanced Msn2 activity. However, we note that nuclear Msn2 is thought to be destabilized during conditions of chronic stress and low PKA activity, as well as heat shock^[32, 35]^.

**Figure 3.**
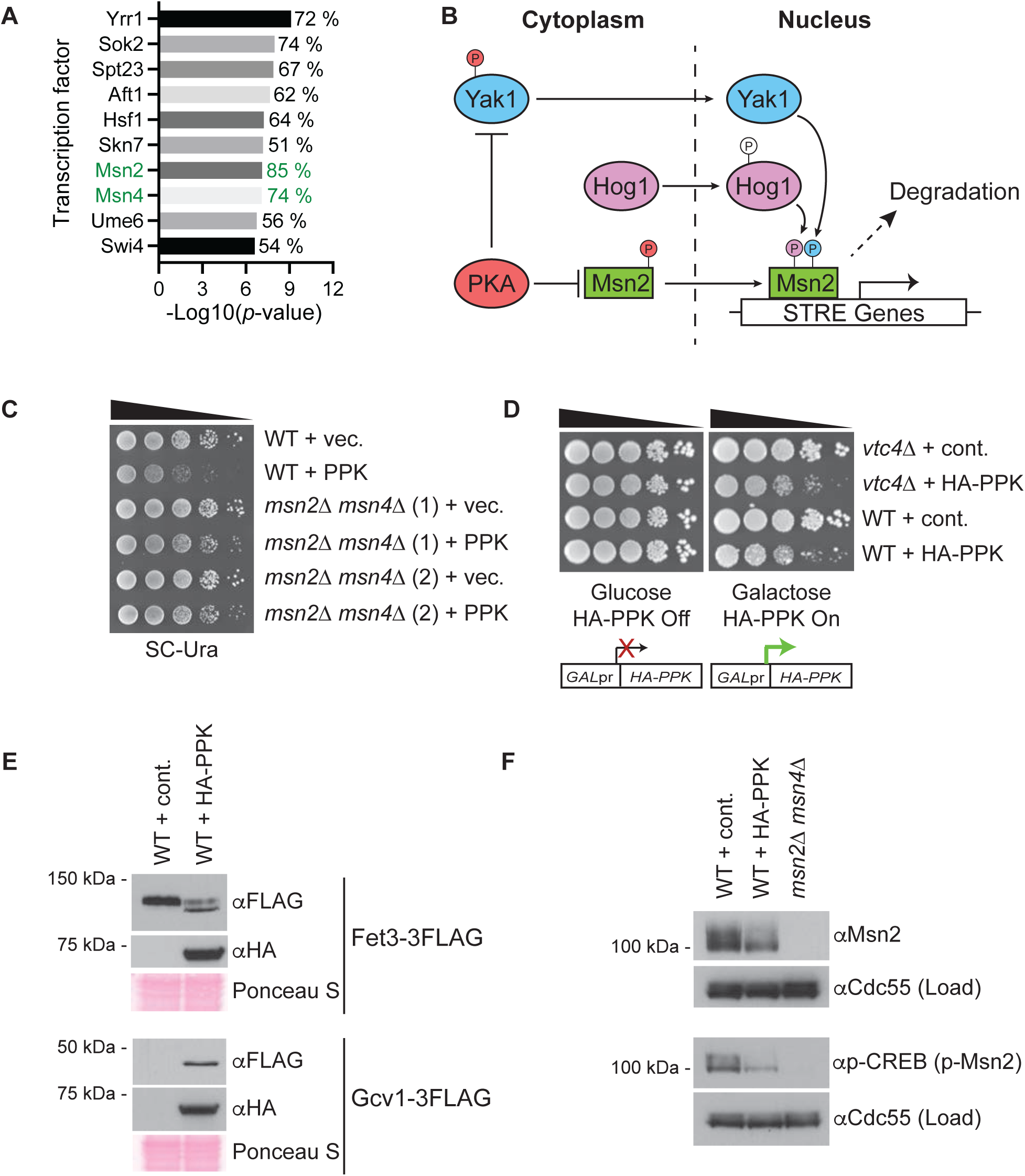
Msn2 and Msn4 transcription factors limit viability in PPK-expressing cells. **A)** YEASTRACT analysis was used to identify candidate transcriptional regulators based on proteins up- and downregulated in mass spectrometry analysis See Materials and Methods for details. The % numbers show the fraction of misregulated proteins (adjusted *p*-value ≤ 0.1) whose encoding genes are regulated by the indicated transcription factors. **B)** Relationship between Yak1/Hog1 and Msn2/4 in stress signaling. **C)** 10-fold serial dilutions of the indicated strains transformed with empty or PPK-expressing vector were spotted on the indicated media and incubated for 48 hours at 30 °C. **D)** Schematic of an inducible HA-PPK expression system under control of the GAL-promoter. GAL-induced HA-PPK expression results in decreased growth in both *vtc4*Δ and wild-type background. 10-fold serial dilutions of the indicated strains were spotted on the indicated media and incubated for 72 hours at 30 °C. **E)** Protein extracts from the indicated strains harbouring the inducible HA-PPK expression system were prepared using a TCA-lysis protocol and separated on SDS-PAGE gels. Separated proteins were transferred to PVDF membrane prior to detection of proteins with the indicated antibodies. Ponceau S stain of the PVDF membranes is shown as loading controls. **F)** Protein extracts from the indicated strains were prepared and analyzed as described in (E). A phospho-CREB antibody was used to detect the NLS specific phosphorylation of Msn2. Cdc55 is used as a loading control. Images representative of n ≥ 3 experiments.

Hog1 and Yak1 are upstream kinases that act on Msn2 to regulate its nuclear import, and at the promoter level to stimulate stress-dependent gene induction (**Fig 3B**)^[36, 37]^. Therefore, we next tested if Hog1 and Yak1 might be involved in the response to HA-PPK expression and polyP accumulation outside of the vacuole. While we saw no change in the expression of Hog1-GFP or Yak1-GFP (**Fig 4A and 4B**), the electrophoretic mobility of Yak1-GFP was slightly but reproducibly increased with PPK expression (**Fig. 4B**). We surmised that this could represent a loss of autophosphorylation or PKA-dependent phosphorylation that accompanies Yak1 activation or translocation to the nucleus, respectively^[38]^. Indeed, PKA activity shows broad mis-regulation in PPK-expressing cells (**Fig. 4C**). Importantly, deletion of *HOG1* or *YAK1* partially rescued the slow growth associated with PPK expression (**Fig. 4D**), and a double *hog1*Δ *yak1*Δ mutant had an even more dramatic effect (**Fig. 4D**). As a control, these strains had equal levels of HA-PPK expression (**Supplementary Fig. 3**). Collectively, our data suggest that Hog1 and Yak1 may signal through Msn2/Msn4 in response to polyP accumulation outside of the vacuole, and that activation of this signaling pathway inhibits growth.

**Figure 4.**
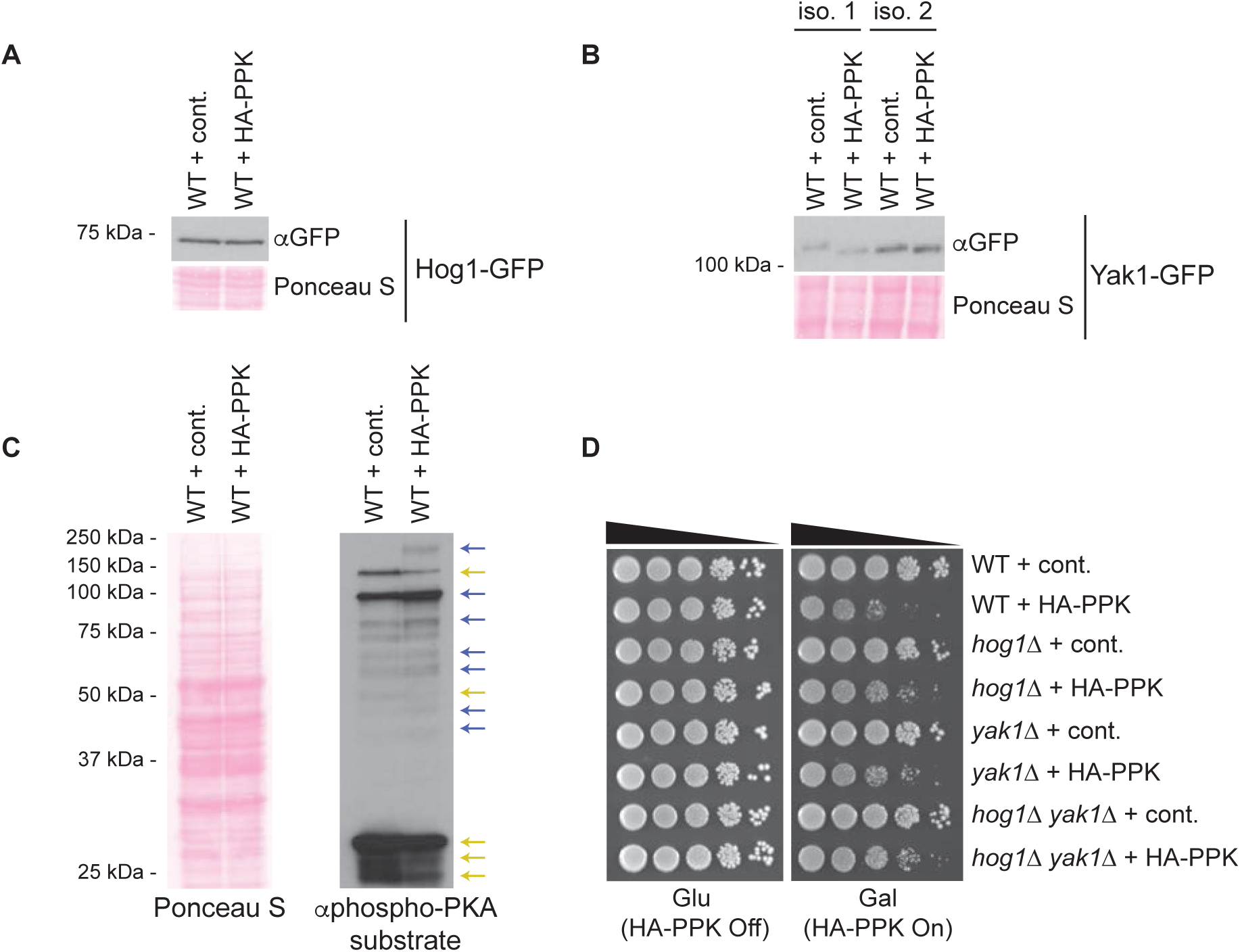
Yak1 and Hog1 kinases modulate the response to PPK-expression. **A)** Hog1-GFP levels are unaffected by PPK expression. Protein extracts from the indicated strains were prepared using a TCA-lysis protocol and separated on SDS-PAGE. Separated proteins were transferred to PVDF membrane before probing with indicated antibodies to detect proteins of interest. Ponceau S stain of the PVDF membrane is shown as a loading control. **B)** PPK expression leads to an increase in the electromobility of Yak1. Protein extracts from the indicated strains were prepared and analyzed as described in (A), but proteins were run on a 10 % SDS-PAGE gel with a 77 : 1 acrylamide to bis ratio. **C)** Phosphorylated PKA substrates were detected using an antibody that detects phosphorylated serine or threonine residues with arginine at the -3 and -2 positions (RRXS*/T*). Blue arrows indicate upregulated phosphorylated PKA substrates and yellow arrows indicate downregulated substrates. Ponceau S stain of the PVDF membrane is shown as a loading control. **D)** 10-fold serial dilutions of the indicated strains were spotted on the indicated media and incubated for **72** hours at 30 °C. Images representative of n ≥ 3 experiments. **Also see Supplementary Figure 3.**

### Ddp1 cooperates with Ppx1 to degrade non-vacuolar polyP

Since aberrant polyP outside of the vacuole leads to changes in expression of the proteome, we expect that mechanisms may exist to counter its accumulation. In yeast, work on polyphosphatase enzymes has focused on four proteins: Ppn1, Ppn2, Ppx1 and Ddp1. Ppn2 and Ddp1 are thought to act as endopolyphosphatases, which cleave polyP chains internally^[2]^. Ppx1 functions as an exopolyphosphatase that cleaves polyP chains starting at the end^[2]^. Ppn1 has been proposed to have both functions^[2, 39]^. Work form the Mayer lab demonstrated that GALpr-*PPX1* strains show decreased growth when grown in glucose (repressed *PPX1*) versus galactose (*PPX1* overexpression) in the presence of non-vacuolar polyP^[4]^. Whether Ppx1 works alone or in conjunction with additional polyphosphatases is unknown. Therefore, we tested the contribution of each of the four polyphosphatase enzymes in our system. Deletion of polyphosphatases *PPN1* or *PPN2* had no impact on the growth or viability of HA-PPK-expressing cells (**Fig. 5A**). This observation is consistent with their roles as vacuolar enzymes and the lack of impact of vacuolar polyP stores on PPK-induced toxicity (**Fig 3D**). In contrast, and as expected, *PPX1* deletion greatly enhanced the toxicity associated with PPK expression (**Fig. 5B**). Interestingly, a similar phenotype was observed in cells disrupted for *DDP1*. In contrast to the slow growth phenotype conferred by PPK expression alone, deletion of these genes resulted in a dramatic decrease in viability in addition to slow growth (**Fig. 5B**). Interestingly, simultaneous deletion of *DDP1* and *PPX1* showed an effect that was not more dramatic than that observed in either single mutant (**Fig. 5B**). This largely epistatic relationship was recapitulated in the context of a liquid growth assay (**Fig. 5C**), suggesting that Ddp1 and Ppx1 function in the same pathway to counter the toxicity of polyP outside of the vacuole. Interpretation of these data is complicated somewhat by our observation that HA-PPK expression is lower in the double *ppx1*Δ *ddp1*Δ relative to wild-type cells or either single mutant (**Fig. 5G and Supplementary Fig. 4A**). To test if Ddp1 and Ppx1 physically interact to degrade polyP in a concerted fashion, we attempted to co-immunoprecipitate 3HA-Ddp1 and Ppx1 but did not detect any interaction between the two proteins (**Supplementary Fig. 4B**).

**Figure 5.**
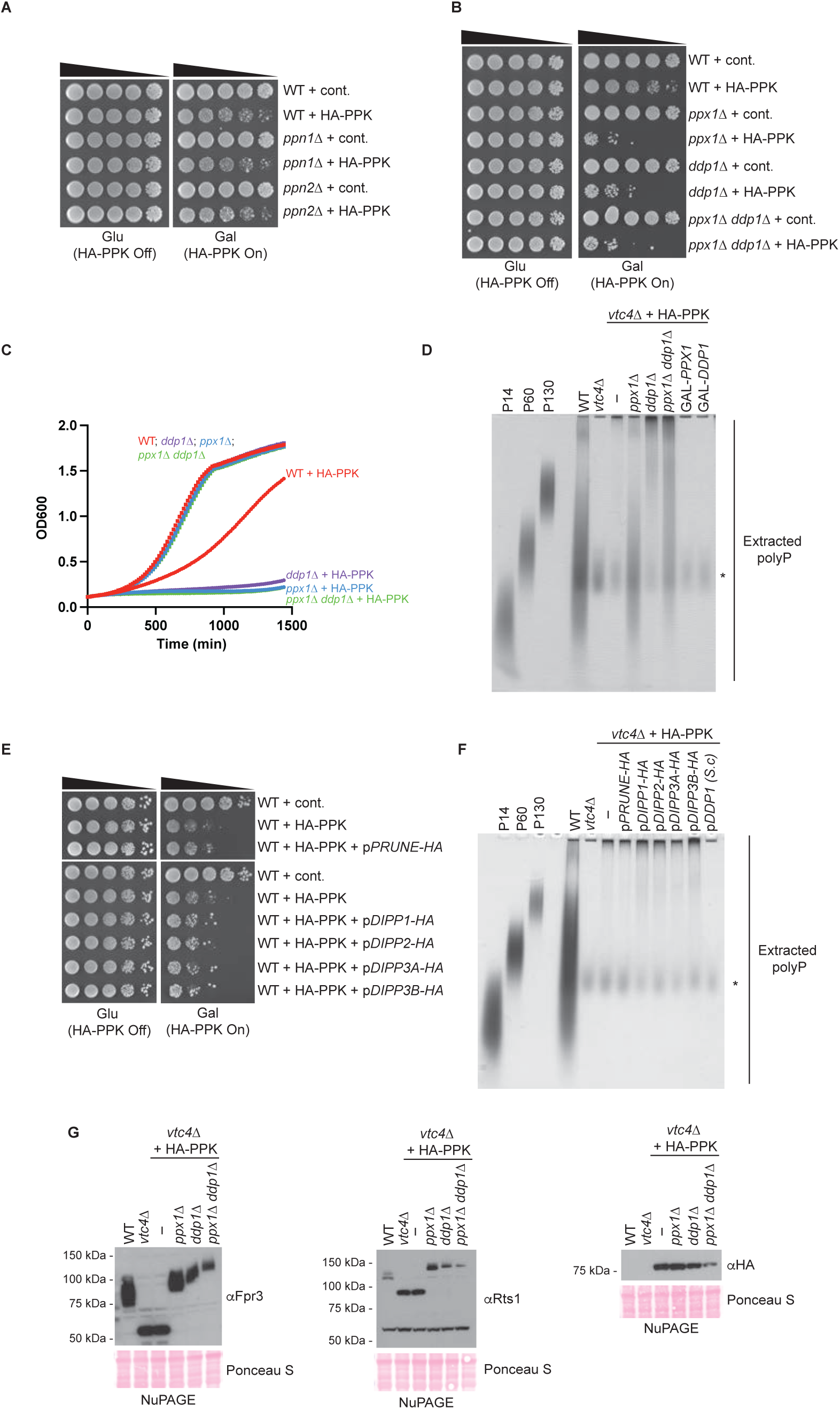
PolyP accumulation outside of the vacuole is countered by Ppx1 and Ddp1. **A-B)** The indicated strains were spotted in a 5-fold dilution series on plates containing 2 % glucose or 1.8 % raffinose + 0.2 % galactose and incubated at 30 °C for 72 hours. **C)** Growth curves for the indicated strains were generated using the Bioscreen C system described in the materials and methods. **D)** Polyphosphate was extracted from the indicated strains and visualized using toluidine blue staining following separation on a 15.8 % acrylamide TBE-Urea gel. The asterisk indicates non-specific staining of an unknown product that is not polyP. **E)** Strains with the indicated plasmids expressing yeast or candidate human polyphosphatase enzymes were spotted in 10-fold dilution series and incubated at 30 °C for 72 hours. **F)** Polyphosphate was extracted from the strains expressing the indicated plasmids and analyzed as in (D). **G)** After a 1 hour galactose induction of HA-PPK, lysine polyphosphorylation of Fpr3 and Rts1 was analyzed by NuPAGE followed by transfer to PVDF membrane and detection of proteins using the indicated antibodies. Ponceau S stain of the membrane is used as a loading control. Images representative of n ≥ 3 experiments. **Also see Supplementary Figures 4 and 5.**

To understand the mechanisms of Ppx1 and Ddp1 action, we analyzed polyP from our panel of polyphosphatase mutants. These experiments were conducted in a *vtc4*Δ background to allow visualization of HA-PPK-synthesized polyP rather than the VTC-synthesized polyP stored in the vacuole but released during the polyP isolation protocol. As we observed in the plasmid-based system, HA-PPK-synthesized polyP accumulated as predominantly long chains reminiscent of those synthesized by *E. coli* in response to cellular stress (**Fig. 5D**). Here again, the amount of polyP is far below that of a wild-type yeast cell, consistent with the notion that it is the localization of HA-PPK-synthesized polyP rather than the amount of that polyP that drives changes in cell growth^[4]^. HA-PPK-expressing strains lacking *PPX1* had more polyP, and the chains were of a medium length (**Fig. 5D**). Strains lacking *DDP1* also showed increased polyP, but these chains were of the long form (**Fig. 5D**). The *ppx1*Δ *ddp1*Δ double mutant accumulated polyP chains of both medium and long lengths, with long chains being predominate. Overexpression of either enzyme reduced the level of PPK-synthesised polyP to background levels (**Fig. 5D**). While these experiments suggest that Ddp1 and Ppx1 act sequentially to degrade long chains of polyP, Ppx1 may also target a distinct population of medium-length chains synthesized by HA-PPK.

Yeast Ppx1 and Ddp1 show homology to human Prune and DIPP1-DIPP3, respectively^[40, 41]^. Prune and DIPP1-DIPP3 have exo- or endopolyphosphatase activity *in vitro*^[40–42]^. Only DIPP1 (also called NUDT3) has been described to have activity *in vivo*^[42]^. Samper-Martin *et al.* show that DIPP1 overexpression confers a zinc-dependent reduction in polyP detected in HEK293T cells via a polyP binding probe^[42]^. Conversely, DIPP1 knockdown caused an increase in polyP in both HEK293T and SH-SY5Y cells using this same method of detection^[42]^. To determine if candidate polyphosphatases could impact PPK-induced toxicity and/or polyP accumulation in yeast, we expressed them in yeast under the strong GPD promoter (**Supplementary Fig. 5A**). In contrast to *DDP1* overexpression, ectopic expression of human Prune or DIPP1-DIPP3 proteins did not improve growth in PPK-expressing cells (**Fig. 5E**) and did not decrease polyP accumulation (**Fig. 5F**). Since DIPP1 activity is thought to be zinc dependent^[42]^, we added zinc to growth media used for these experiments. Although the highest concentrations of zinc used caused slight improvement of growth of strains expressing HA-PPK alone or in the presence of Prune, growth of DIPP1-DIPP3 expressing strains was unaffected (**Supplementary Fig. 5B**). Similarly, although the addition of zinc resulted in a slight decrease in polyP levels in all strains tested, this was not impacted by the ectopic expression of DIPP1 (**Supplementary Fig 5C**). Either Prune and DIPP1-DIPP3 do not directly impact the accumulation of long chain polyP *in vivo*, or their activities are not faithfully recapitulated in yeast under the conditions tested.

To corroborate and extend our findings with yeast Ppx1 and Ddp1, we analyzed lysine polyphosphorylation of previously identified targets^[14]^. Since lysine polyphosphorylation is a non-enzymatic modification, it is thought to be controlled largely by the concentration of polyP itself. We carried out these experiments in a *vtc4*Δ background to prevent interference of vacuolar polyP which can polyphosphorylate proteins during cell lysis^[43]^. Deletion of *PPX1* or *DDP1* in PPK-expressing cells resulted in decreased electrophoretic mobility of polyphosphorylation targets Rts1 and Fpr3 on NuPAGE gels, indicative of lysine polyphosphorylation (**Fig. 5G**)^[14]^. Therefore, the ‘extra’ polyP in these cells is competent to participate in lysine polyphosphorylation reactions. The decreased electrophoretic mobility observed in *ppx1*Δ and *ddp1*Δ cells could result from the modification of additional lysines at higher concentrations of polyP^[44]^. The greater impact of *ddp1*Δ may reflect the addition of longer chains to individual target lysines in these cells. We propose that the enhanced toxicity of PPK-expression observed in *ddp1*Δ and *ppx1*Δ mutants may be due in part to aberrant polyphosphorylation of proteins in the cytoplasm.

## DISCUSSION

### Non-vacuolar polyP reprograms the yeast proteome

There are conflicting data concerning the concentration of polyP that accumulates outside of the vacuole in wild-type yeast, although it is generally accepted that the vacuole harbours substantially higher concentrations than other areas of the cell^[2]^. The sequestration of polyphosphate in specific subcompartments is not unique to yeast. Human cells accumulate high concentrations of polyP in the nucleolus or in dense granules of platelets^[8, 10]^. Trypanosomes accumulate high levels of polyP in acidocalcisomes^[45]^, as well as glycosomes and the nucleolus in smaller amounts^[46]^. Even in many bacteria, polyP produced in response to stress is concentrated in granules^[2]^. While the concentration of polyP in these areas is important for specific cellular functions (i.e. blood clotting in the case of dense granules^[47]^), it is less clear if polyP is actively excluded from (or limited within) some areas of the cell and why this might be important. Expression of bacterial PPK in yeast, coincident with production of polyP outside the vacuole, was previously demonstrated to reduce growth and sensitize cells to rapamycin and cycloheximide^[4, 14]^. Our new study provides a critical advance by *i)* identifying molecular pathways that mediate toxic effects of PPK expression, and *ii)* providing molecular insight into how exo- and endopolyphosphatases cooperate to counter the accumulation of non-vacuolar polyP in yeast.

We are not able to rule out that other enzymatic activities of the PPK enzyme (e.g. phosphorylation of nucleotides), contribute to the proteomic changes measured in our mass spectrometry experiments. However, the fact that PPK’s impact is greatly exacerbated in strains lacking the Ddp1 and/or Ppx1 polyphosphatases, suggests that downstream changes in growth and/or viability are related to changes in polyP metabolism. PolyP could influence the expression or stability of some proteins directly, perhaps by binding to transcription factors or proteins themselves. We expect the investigation of such direct modes of action will provide clues to molecular functions of polyP that are broadly applicable, even under endogenous scenarios. Protein changes following PPK expression could also be indirect, for example, as a downstream result of polyP-mediated ion chelation or the cell’s response to more direct activities.

Importantly, we consider such indirect effects to be equally interesting, as they provide valuable insights into how eukaryotic cells respond to mislocalized polyP and may aid in identification of scenarios when this occurs in the absence of PPK expression.

### Ddp1 and Ppx1 limit the toxic effects of non-vacuolar polyP

In *ppx1*Δ and *ddp1*Δ *ppx1*Δ cells expressing PPK, polyP accumulates as medium-length chains, similar to those made by the VTC complex in wild-type cells. This argues against the idea that PPK expression is toxic only because it makes long chains that are not normally seen in yeast. However, we can’t rule out that different chain lengths have unique impacts on cell signaling and physiology and this remains an important area for future research. Indeed, there is precedent for chain-length dependent effects of polyP. For example, polyPs of different length affect different parts of the blood coagulation cascade^[48]^, and long chains preferentially stimulate changes in macrophage signaling important for the immune response^[49]^. Since we have not recovered any evidence of a physical interaction between Ddp1 and Ppx1, we propose that they act sequentially to degrade long polyP chains (**Fig. 6**), with Ddp1 first cleaving chains to generate free polyP ‘ends’ on which Ppx1 can then act. Notably, Ppx1 has also been shown to directly remove polyP chains from nuclear polyphosphorylated target Nsr1^[50]^, and could act on other proteins in PPK-expressing cells to limit toxicity.

**Figure 6.**
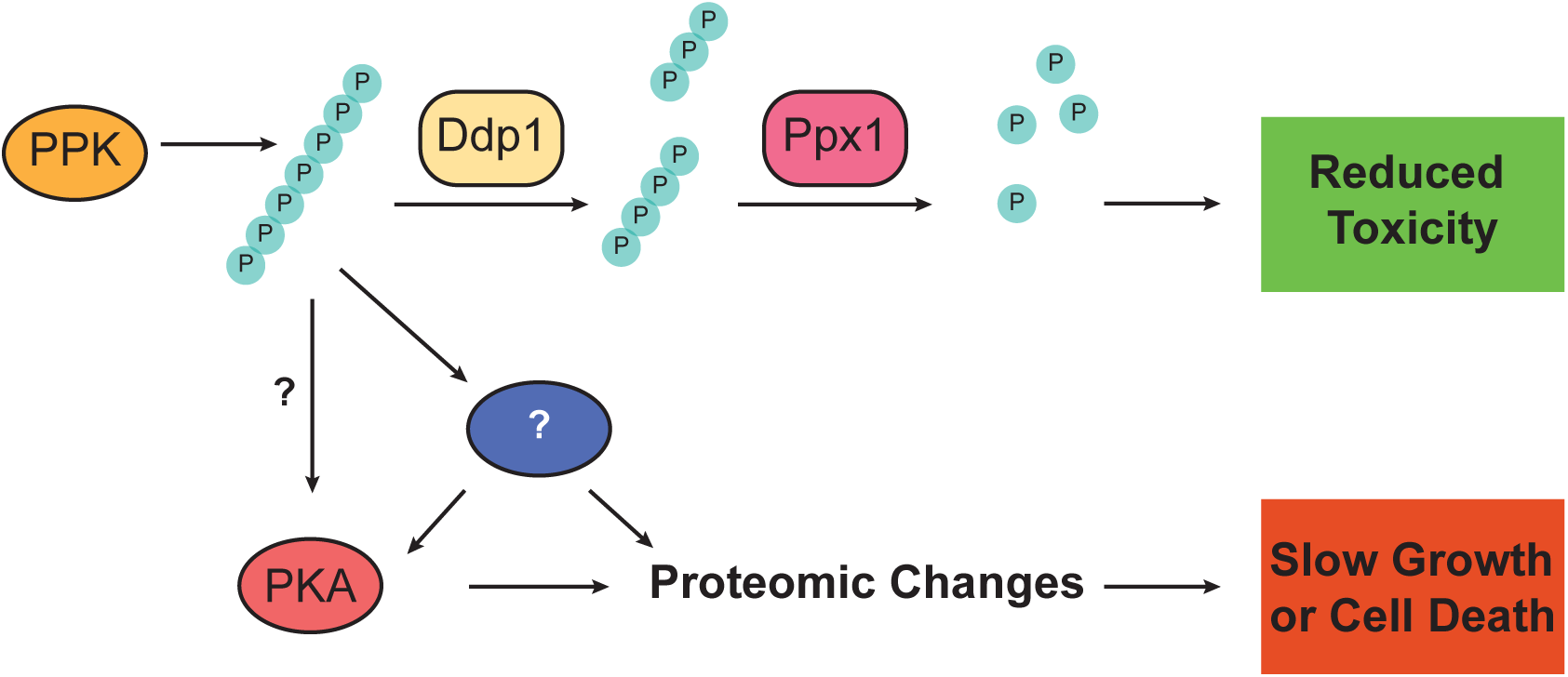
Working model for the cellular response to non-vacuolar polyP in yeast. PPK- produced polyP accumulates in the cytoplasm and slows growth by a variety of pathways which include activation a Yak1/Hog1-Msn2/4 signaling cascade. The accumulation of polyP outside of the vacuole is countered by Ppx1 and Ddp1 which work together to degrade polyP. See text for details.

### PolyP as an activator of stress signaling

We propose that polyP escaping degradation by polyphosphatases activates a stress-signaling pathway that limits cell growth or viability (**Fig. 6**). It is unclear if polyP is indeed causing some type of intracellular stress (i.e. accumulation of unfolded proteins), or if it is inappropriately activating stress-signaling pathways in the absence of a stress. Regardless, the downstream consequences appear to involve the PKA pathway, which plays a central role in the regulation of cell metabolism, growth, as well as diverse stress responses^[51]^. Our results show that accumulation of non-vacuolar polyP leads to broad misregulation of PKA substrates. Some of these substrates may include misregulated proteins identified in our mass spectrometry experiments. For example, Ura2, which is downregulated in PPK-expressing cells, is phosphorylated by PKA *in vitro*^[52]^. Moreover, PKA regulates the activity of Msn2/4 and Yak1 through direct phosphorylation^[33, 38]^, and recently it has been suggested that PKA might modulate the activation of the Hog1 pathway during impaired sphingolipid biosynthesis^[53]^. Previous work has shown that deletion of *YAK1* or *MSN2* and *MSN4* under conditions of low PKA activity rescues related slow-growth phenotypes^[54, 55]^. We have shown that deletion of *YAK1* or *MSN2* and *MSN4* also rescues growth in PPK-expressing cells, which is in accordance with scenarios of low PKA activity. Further, low PKA activity imparted by the accumulation of non-vacuolar polyP could explain the sensitivity of PPK-expressing cells to rapamycin^[14]^, since mutants with low PKA activity have increased sensitivity to this drug^[56]^. Our results suggest that non-vacuolar polyP might affect PKA activity directly or indirectly and that these changes may in part explain the observed toxicity (**Fig. 6**).

### Endogenous functions of non-vacuolar polyP

Polyphosphate functions outside of the vacuole are poorly understood. The Ppx1 and Ppn1 polyphosphatases localize in part to the mitochondria and polyP here is suggested to play a role in energy homeostasis and metabolism^[57]^. Within the nucleus, polyphosphorylation may also inhibit the interaction of Top1 and Nsr1 proteins in addition to modulating their localization to the nucleolus^[50]^. It is also tempting to speculate that nuclear localization of polyphosphate underlies its poorly understood role in cell cycle regulation^[58]^. How polyP that functions in or transits through the cytoplasm in a wild-type cell avoids degradation by polyphosphatases is an intriguing question. It may be that these polyP molecules are bound by proteins or small molecules that prevent enzyme access to the phosphoanhydride bonds. It is also possible that Ddp1 and Ppx1 become activated to degrade cytoplasmic polyP only when the concentration exceeds a particular threshold. We further speculate that polyP could be released from vacuoles following damage with lysosomaltropic agents such as *N*-Dodecylimidazole^[59]^, or during sporulation, where the vacuole undergoes programed lysis^[60]^. Ddp1 and Ppx1 are ideally placed to regulating these responses.

### Regulation of polyP in human cells

We previously found that transient expression of PPK in human cells also resulted in polyP accumulation throughout the cell^[61]^. Accumulation of long-chain polyP was associated with broad changes in gene and protein expression, increased phosphorylation of ERK1/2 and p70S6K, and redistribution of multiple nuclear proteins from the nucleus/cytoskeletal fraction to the cytoplasm and/or organelles^[61]^. We did not observe activation of apoptotic pathways or signs of extensive cell death^[61]^, consistent with a previous report^[62]^. However, it is possible that in the context of our transient expression system, plasmid copy number, PPK expression, and/or polyP itself are quickly downregulated to manageable levels. Based on our work in yeast, we expect that inhibition of polyphosphatases will increase levels of polyP to levels that confer greater toxicity. Our study further suggests that degradation of long-chain polyP in human cells will require the combined action of both endo- and exopolyphosphatases, akin to what we observed for Ddp1 and Ppx1. Mammalian polyphosphatases may also play a role in degrading long-chain polyP made by pathogenic bacteria, which has been suggested to enter cells during infection to reprogram macrophages and thereby subvert host defense mechanisms^[49]^. The identities of intracellular human polyphosphatases had long remained unknown, but Prune and DIPP1-DIPP3 have emerged as candidates based on their *in vitro* activities^[40,41,63]^. Notably, mutations that reduce Prune activity *in vitro* give rise to the rare disease called <u>N</u>eurodevelopmental disorder with <u>mi</u>crocephaly, <u>h</u>ypotonia and variable <u>b</u>rain <u>a</u>nomalies (NMIHBA)^[63]^. However, expressing either Prune or DIPP1-DIPP3 in yeast resulted in no appreciable change in viability and no obvious decrease in PPK-synthesized polyP. We note that the *in vitro* exopolyphosphatase activity of Prune is mostly against very short chains (3-4 phosphate units in length)^[41, 63]^. In our experiments, Prune was also expressed at substantially lower levels than the DIPP1-DIPP3 proteins in yeast, despite the fact that both constructs used the GPD promoter. Together, these observations could account for the lack of effect observed in our system. More surprising is the observation that expression of DIPP1 had no effect in either the presence or absence of zinc. This finding is seemingly at odds with the exciting work of Samper-Martin *et al.*, which describes a zinc-dependent activity of DIPP1 (also called NUDT3) in HEK293 and SH-S5Y cells^[42]^. One exciting possibility is that DIPP1 or other mammalian polyphosphatase enzymes require additional co-factors besides zinc that are not normally present in yeast. Such proteins could serve as important regulators of *in vivo* polyphosphatase activity and should be a focus of future work.

## MATERIALS & METHODS

### Yeast strains & handling

Yeast strains were constructed as previously described using standard methods^[64–66]^. Locations of gene deletion and tagging cassettes were confirmed using PCR analyses^[66]^. Full genotypes of all yeast strains used in this work are listed in **Supplementary Table S2.** Cells were grown in YPD, YEP-Raf. YEP-Gal, or synthetic complete media as described below for individual experiments. In general, SC-Ura and SC-Leu media were used to maintain *URA3*-based and *LEU2*-based selection plasmids, respectively.

### Plasmids

All plasmids used for this work are listed in **Supplementary Table S5.** All plasmids generated for this work will be made available from Addgene (www.Addgene.org) following publication.

### Proteomics

The mass spectrometry proteomics data have been deposited to the ProteomeXchange Consortium via the PRIDE (PMID: 34723319) partner repository with the dataset identifier PXD031326. Reviewers can access the data in PRIDE using the following username: and password: oZF6dkud. Summary data is provided in **Supplementary Table S1**.

### Polyphosphate extractions

#### Cell growth

Cells expressing indicated plasmids in **Fig. 1A** and **Fig. 2C** were diluted to OD600 = 0.2 in SC-Ura + 2 % glucose and grown at 30 °C with shaking until the OD600 reading reached a final value of 0.8-1. Cells **in Fig. 5D** were diluted to OD600 = 0.2 in YEP + 2 % raffinose and grown for 3 hours before adding 2 % galactose and growing another 3 hours. Cells expressing indicated plasmids in Fig. 5F were diluted to OD600 = 0.2 in SC-Ura + 2 % raffinose and grown for 3 hours before adding 2 % galactose and growing another 3 hours.

#### PolyP extraction

The polyP isolation protocol was adapted from a protocol first described by Bru *et al.* 2017^[67]^ and described elsewhere^[61]^. The protocol is repeated here using similar language. Yeast cell pellets of approximately 8 OD600 units in size were resuspended in 400 µL of cold LETS buffer (100 mM LiCl, 10 mM EDTA, 10 mM Tris-HCl pH 7.4, 0.2 % SDS), transferred to a screw cap tube containing 600 µL of phenol pH = 8 and 150 µL of mqH_2_0, and vortexed for 20 sec. Samples were then heated for 5 min at 65 °C and cooled on ice for 1 minute. Once cooled, 600 µL of chloroform was added and the samples were vortexed for 20 seconds. Vortexed samples were centrifuged for 2 min at 13 000 x g at room-temperature and the top layer was transferred to a new tube containing 600 µL of chloroform. Samples were vortexed and centrifuged again for 2 min at 13 000 x g, and the top layer was transferred to a clean 1.5 mL tube. To degrade RNA and DNA, 2 µL of 10 mg/mL RNAse A and 2 µL of 10 mg/mL DNAse I were added to samples and incubated at 37 °C for 1 hour. Digested samples were transferred to pre-chilled 1.5 ml tubes containing 1 ml of 100 % ethanol and 40 µl of 3 M sodium acetate pH = 5.3 and polyP was left to precipitate at -20 °C for at least 3 hours or overnight. PolyP was pelleted by centrifugation for 20 min at 13 000 x g at 4 °C and the supernatant was discarded. The pellet was washed with 500 µL of 70 % ethanol and centrifuged for an additional 5 min. The supernatant was discarded and the pellets were air dried before being resuspended in 20-30 µL of mqH_2_0. *Gel analysis:* Extracted polyP was diluted 1:1 with polyP loading buffer (10 mM Tris-HCl pH 7, 1 mM EDTA, 30 % glycerol, bromophenol blue) and 15 µL was loaded on a 15.8 % TBE-Urea gel (5.25 g Urea, 7.9 mL 30 % acrylamide, 3 mL 5 X TBE, 150 μL 10 % APS, 15 μL TEMED) and run at 100 V for 1 h 45 min in 1 X TBE running buffer. For toluidine blue staining, the gel was incubated in fixing solution with toluidine blue (25 % methanol, 5 % glycerol, 0.05 % toluidine blue) for 20 min and destained with fixative solution without toluidine blue. For DAPI staining, the protocol was adapted from Smith and Morrisey 2007^[68]^. Extracted polyP was diluted 1 : 60 for wild-type strains and 1 : 10 for *vtc4*Δ with polyP loading buffer and 15 µL was loaded. The gel was incubated in fixing solution with DAPI (25 % methanol, 5 % glycerol, 50 mM tris base, 2 ug/mL DAPI) for 30 minutes. The gel was destained with fixative solution without DAPI for 1 hour, changing solution once halfway through. The destained gel was exposed to 365 nM UV light on a transilluminator to photobleach the DAPI-polyP until it became dark. Pictures were taken with a Bio-Rad ChemiDoc UV transilluminator.

### Label-free mass spectrometry

#### Sample preparation

Sample preparation and mass spectrometry protocols are based on an experimental set up we have used previously for work in HEK293T cells^[61]^. Similar language is used in the description here. Cells were diluted to OD600 = 0.2 in SC-Ura with 2 % glucose and grown at 30 °C with shaking until OD600 = 1. Yeast pellets of approximately 50 OD600 units in size were resuspended in 600 µL of lysis buffer (5% SDS, 50mM TEAB, supplemented with protease and phosphatase inhibitor tablets (Roche)) and the 2 mL screwcap tube was filled to capacity with acid-washed glass beads. Cells were lysed with 12 x 1 minute pulses on a BioSpec mini bead-beater with incubation on ice in between pulses. Lysates were clarified by centrifugation at 15 000 x g for 10 min at 4 °C. The supernatant was collected and centrifuged again at 15 000 x g for 10 min at 4 °C before being collected again and frozen at -80 °C. Frozen protein extracts were shipped overnight on dry ice to the UC Davis Proteomics core. Trypsin digestion was performed using S-Trap Mini spin columns (PROTIFI) as per the manufacturer’s protocol.

#### Liquid chromatography and mass spectrometry analysis

Digested peptides were first separated using Proxeon Easy-nLC II HPLC (Thermo Scientific) and Proxeon nanospray source. Digested peptides were loaded on a 100 micron x 25 mm Magic C18 100 Å 5U reverse phase trap where they were desalted on-line before being separated using a 75 micron x 150 mm Magic C18 200Å 3U reverse phase column. Peptides were eluted using a 140 minute gradient with a flow rate of 300 nl/min into a Thermo Scientific Q Exactive Plus Orbitrap mass spectrometer. An MS survey scan was obtained for the m/z range of 350-1600. MS/MS spectra were acquired with the top 15 ions in the MS spectra subjected to Higher Energy Collisional Dissociation, with a resolution at 400 m/z of 70,000 and 17,500 at the MS1 and MS2 levels, respectively. For precursor ion selection, an isolation mass window of 1.6 m/z was used. A normalized collision energy of 27% was used for fragmentation. Dynamic exclusion was set for fifteen seconds.

### Bioinformatics analyses

For mass spectrometry data, the bioinformatics pipeline used for analysis is similar to that previously described^[61]^. The details are presented here with minimal rewording for clarity.

### Protein identification

Tandem mass spectra were extracted using the msConvert program^[69]^. X!Tandem (the gmp.org; version X! Tandem ALANINE (2017.2.1.4)) was used to analyze MS/MS spectra and identify peptide sequences^[70]^. The X!Tandem program was used to search the SwissProt Uniprot *S. cerevisiae* database^[71]^ (proteome UP000002311, date of download = 2018-12-21), along with 110 common lab contaminants (thegpm.org/crap/) and the *E. coli* PPK protein sequence, and all of these sequences reversed for a decoy database search. A database search was performed on MS/MS spectra from precursor ions with a charge state of at most 4. Trypsin was set as the digestion enzyme. Using X!Tandem two-pass search, the initial search was performed with one miscleavage considered, and a refinement search looked for additional unanticipated miscleavages in peptides from proteins identified in the first pass. Fragment ion and parent ion mass tolerances were set to 20 ppm. Single amino acid polymorphisms were checked for each peptide residue in the first search, using a default list of polymorphisms from the SNP annotations in ENSEMBL^[72]^. Carbamidomethylation of cysteine and selenocysteine were specified as fixed modifications. Variable modifications included were: glutamic acid->pyro-glutamic acid of the N-terminus, glutamine->pyro-glutamine of the N-terminus, ammonia loss of the N-terminus, deamidation of asparagine and glutamine, oxidation and dioxidation of both methionine and tryptophan, and N-terminal acetylation. Validation of peptide and protein identifications was done using the Scaffold program (version 4.11.1 from Proteome Software, Inc, Portland OR). The ProteinProphet algorithm was used to assign protein identification probabilities. Peptide identifications associated with an FDR < 1% were deemed of high confidence. Protein identifications with an FDR < 1% and associated with at least 2 high-confidence peptides were then used in all downstream analyses. Proteins were grouped into protein groups if they shared significant peptide evidence. To satisfy the principles of parsimony, proteins sharing the same peptides were grouped if it was not possible to differentiate them based on MS/MS analyses.

### Differential protein expression analysis

Protein and protein group spectral counts were retrieved from Scaffold. To account for missing values, spectral counts were imputed for all spectral count values equal to zero by randomly sampling spectral count values from the bottom 20 % of non-zero spectral counts. Spectral counts were normalized by the total number of spectral counts in each MS/MS runs. Protein differential expression was assessed using a two-tailed, two-sample Student’s *t*-test assuming unequal variance on normalized spectral counts. Differential expression was assessed using a *t*-test only for proteins that were reproducibly identified in 3 or more replicates in both experimental conditions. The Benjamini-Hochberg procedure^[73]^ was used to correct for multiple hypothesis testing. Proteins were deemed differentially expressed when their FDR-adjusted *p*-value was smaller or equal to 0.1. In addition, proteins with extreme expression differences (i.e. detected in all replicates in one condition and in none for the other condition) are also reported and named “all-or-none” proteins.

### Gene Ontology enrichment analysis

In order to investigate key functional changes in response to ectopic expression of PPK1, a Gene Ontology^[74]^ enrichment analysis was performed using Ontologizer^[75]^. Differentially expressed proteins (FDR-adjusted *p*-value ≤ 0.1) and all-or-none proteins were evaluated for enrichment against a background of all identified proteins in all MS/MS runs (FDR < 1%). The OBO ontology file and GAF annotation file used for gene ontology enrichment analysis were downloaded from http://geneontology.org/ on 2019-08-22 and 2020-01-20, respectively. The *p*-values were adjusted using the Benjamini-Hochberg procedure to correct for multiple hypothesis testing.

### Protein-protein interaction network

The STRING^[21]^ database was used to construct a protein-protein interaction network between differentially expressed proteins (FDR-adjusted *p*-value ≤ 0.1) and all-or-none proteins. The network was constructed using the STRING medium confidence (0.4) threshold for experimental evidence and database evidence of interactions. Proteins without any protein-protein interactions were removed from the network.

### YEASTRACT analysis

Genes of differentially expressed proteins (FDR-adjusted *p*-value ≤ 0.1) and all-or-none proteins were ranked by transcription factor using YEASTRACT (http://www.yeastract.com/formrankbytf.php)^[28]^. Documented regulations were filtered by DNA binding or expression evidence, transcription factor acting as activator or inhibitor, and all transcription factors were checked.

### Immunoprecipitation

Cells were diluted to OD600 = 0.2 in YEP + 2 % raffinose and grown for 3 hours before inducing 3HA-Ddp1 expression with 2 % galactose for 2 hours. Yeast cell pellets of approximately 50 OD600 units were resuspended in 750 µL of lysis buffer (0.1 % NP- 40, 150 mM NaCl, 50 mM Tris-HCl pH 8, 10 % glycerol, 15 mL of buffer was supplemented with 30 µL of 1 M imidazole, 15 µL of 1 M sodium fluoride, 17.25 µL of 1 M sodium-molybdate, 75 µL of 200 mM sodium orthovanadate, 75 µL of 1 M DTT, 15 µL 5 mM cantharidin, 150 µL of 2.5 mM (-)-p-Bromotetramisole Oxalate, 75 µL of 1 M glycerol-2-phosphate, 15 µL of 0.1 M PMSF, and a 15 mL Roche cOmplete protease inhibitor tablet with EDTA^[14, 50]^, and the 2 mL screwcap tubes were filled to capacity with acid-washed glass beads. Cells were lysed with 8 x 1.5 minute pulses on a BioSpec mini bead-beater with incubation on ice in between pulses. Lysates were clarified by centrifugation at 15 000 x g for 10 min at 4 °C.

The supernatant was collected and centrifuged again at 15 000 x g for 10 min at 4 °C. Anti-HA antibody was added to each lysate at a dilution of 1/650 and rotated end over end for 1 hour at 4 °C. After incubating, 20 µL of Protein A Magnetic Beads were washed 3 times with 700 µL of lysis buffer and were added to each sample and rotated end over end for 1 hour at 4 °C. Samples were centrifuged briefly to pellet beads and lysate was removed. Beads were washed 3 times with 700 µL of lysis buffer and eluted with sample buffer by boiling for 10 minutes.

### Immunoblotting

Proteins were extracted using a method that was described previously^[14]^. That method is described again here for convenience. Cell pellets of OD600 = 3-6 units (as measured using an Eppendorf BioPhotometer) were resuspended in 300 µL of 20 % TCA and 100 µL of acid-washed glass beads. Cells were lysed with 2 x 3 minute pulses on a BioSpec mini bead-beater. Supernatant was removed and kept. Beads were washed with 300 µL 5 % TCA and this supernatant was combined with the first 300 µL. This mixture was clarified via centrifugation for 4 minutes at 4 °C at 16 000 x g. The resulting pellet was dissolved in 100 µL of SDS-PAGE loading buffer supplemented with 1/10 volume 1.5M Tris-HCl pH 8.8 and 1/10 volume 1M DTT and boiled for 8 minutes. Samples were centrifuged again at 16 000 x g and the supernatant was recovered. Approximately 10-20 µL were loaded per gel (corresponding to about 75 µg of protein). Unless stated otherwise in figure legends, protein samples were run on 10 % SDS-PAGE gels made from a 30 % acrylamide/bis solution, 37.5 : 1. To resolve polyphosphorylation induced shifts, protein samples were run on NuPAGE 4 to 12 % Bis-Tris gels. Buffers used for both gel types were described previously^[14]^. All proteins were transferred to PVDF membranes before probing with antibodies. The antibodies used for this work are detailed in **Supplementary Table S4.**

### Spot tests

Cells from freshly grown plates were diluted in water and spotted on the indicated media and allowed to dry prior to incubation at 30 °C for 2-3 days.

### Liquid growth curves

Overnight cultures grown in YEP with 2 % raffinose were diluted to OD600 = 0.2 in YEP with 2% raffinose and incubated at 30 °C with shaking for 4.5 hours (OD600 = 0.6-0.8). Cultures were subsequently diluted to OD600 = 0.1 (measured using an Eppendorf BioPhotometer) in YEP with 2 % raffinose or YEP with 2 % galactose. 200 µL of cells were pipetted into wells of a 100 well honeycomb plate and the plate was incubated in a Bioscreen C plate reader (Oy Growth Curves) set at 30 °C with continuous shaking. A wavelength of 600 nm was used to measure optical density every 15 minutes for 24 hours.

## ACKNOWLEDGEMENTS

We thank Dr. T. Shiba (RegeneTiss, Japan) for the gift of polyP standards and Andreas Mayer for the original p416-GPD and p416-GPD-PPK plasmid. We thank the UC Davis Proteomics Core Facility for mass spectrometry analysis. We also thank the Rudner Lab for the Cdc55 antibody, as well as the Baetz Lab for the Msn2 and p-PKA substrate antibodies. Work for this project was funded by Canadian Institutes of Health Research (CIHR) Project Grants to MD (PJT-148722) and MD & MLA (PJT-450860) and a Natural Sciences and Engineering Research Council of Canada (NSERC) Discovery Grant to MLA. Trainees in the Downey Lab are supported in part by an Early Researcher Award from the Ontario Ministry of Research, Innovation and Science. LM was supported by an Ontario Graduate Scholarship and a NSERC Canada Graduate Scholarship. IA was supported by an Ontario Graduate Scholarship and a NSERC Canada Graduate Scholarship.

## FIGURE LEGENDS

**Supplementary Figure 1.**
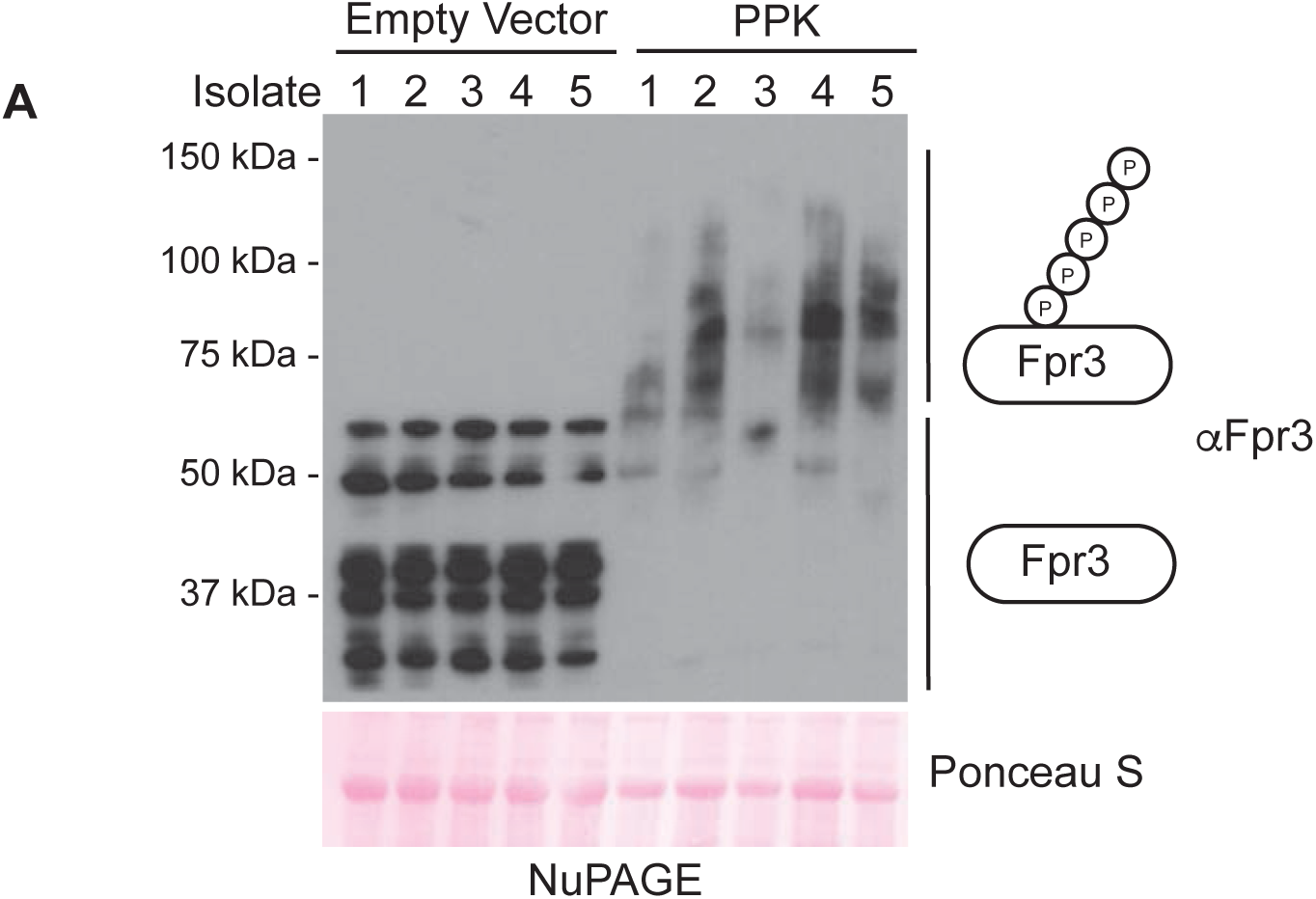
Changes to the *S. cerevisiae* proteome with ectopic PPK expression. PPK expression restores polyphosphate levels in samples used for mass spectrometry. Protein extracts were prepared from the indicated strains carrying either control or PPK expressing plasmids using an SDS lysis protocol and separated on a NuPAGE gel. Separated proteins were transferred to PVDF before probing with an antibody that recognizes yeast Fpr3. Ponceau S stain of the membrane is shown as a loading control. The shift observed in PPK-expressing samples is a sign of lysine polyphosphorylation and polyP production.

**Supplementary Figure 2.**
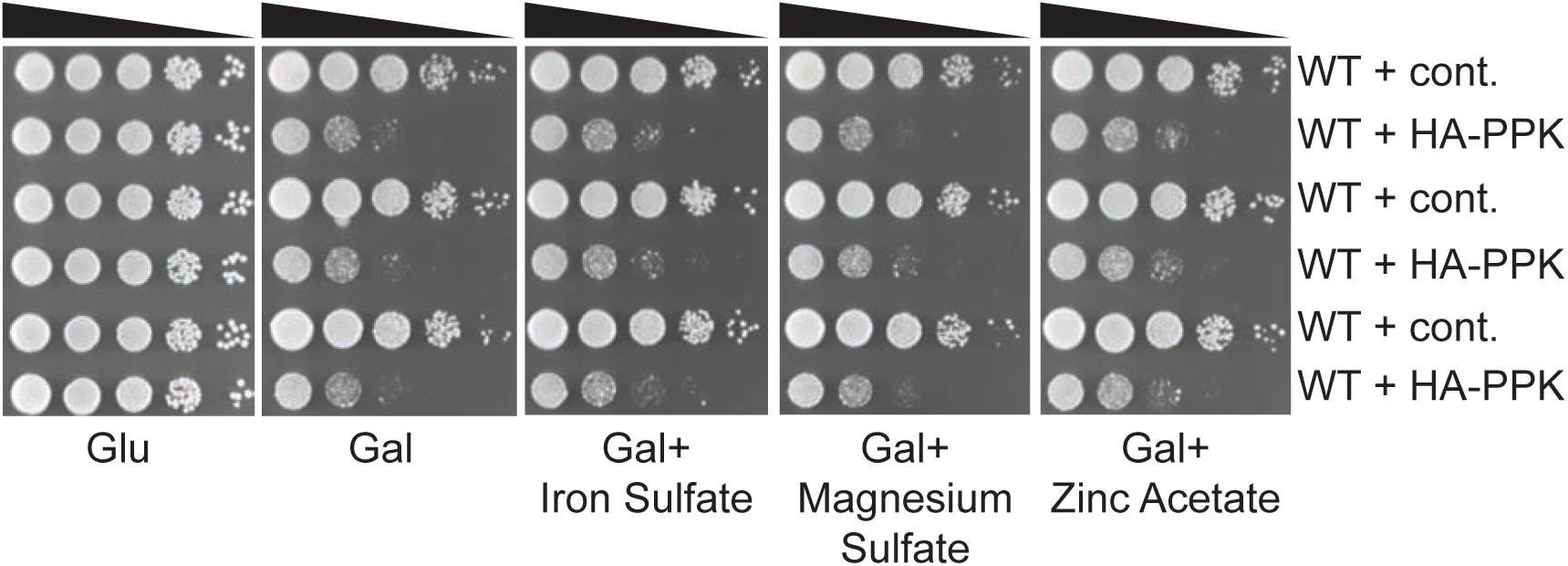
Supplementing media with excess metals does not rescue PPK-induced growth defects. The indicated strains were spotted in 10-fold dilution series on media containing 200 μM of the indicated metals and incubated at 30 °C for 48 hours. Images representative of n = 2 experiments.

**Supplementary Figure 3.**
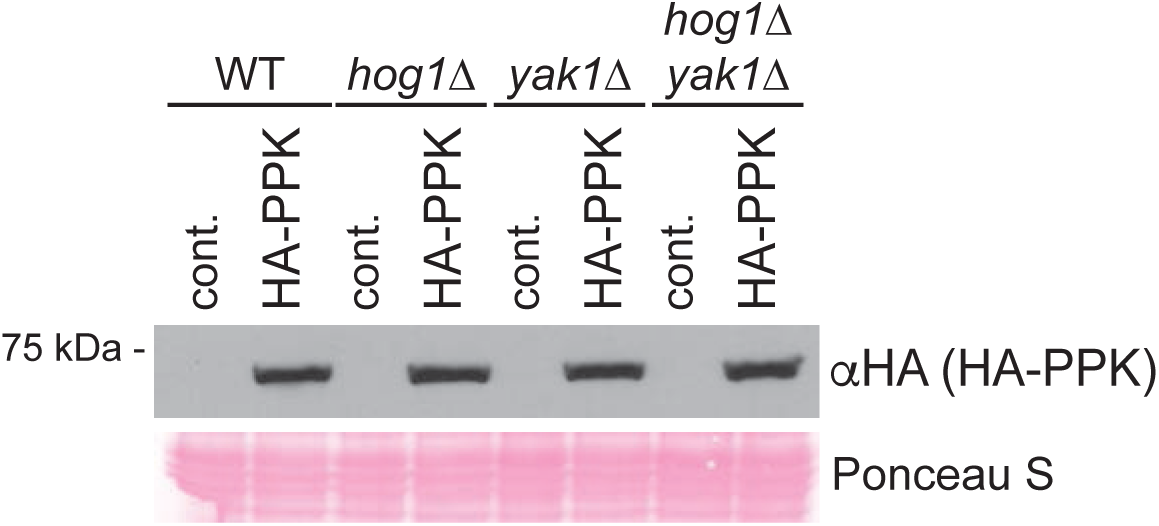
HA-PPK levels in the *HOG1* and *YAK1* mutants. Protein extracts from the indicated strains were prepared using a TCA-lysis protocol and separated on SDS-PAGE. Separated proteins were transferred to PVDF and probed with anti-HA to detect HA-PPK. Ponceaus S stain is shown as a loading control. Images representative of n = 3 experiments.

**Supplementary Figure 4.**
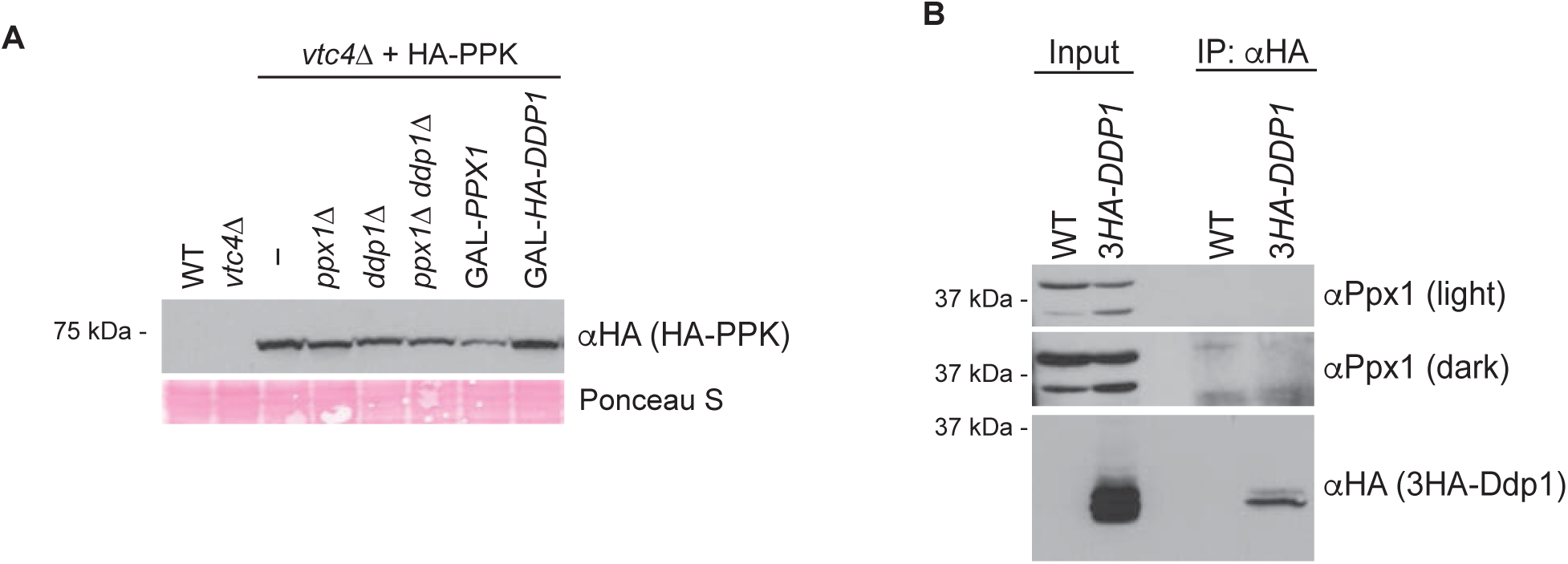
HA-PPK regulation in polyphosphatase mutants. **A)** PPK levels in polyphosphatase mutants. Protein extracts from the indicated strains were prepared using a TCA-lysis protocol and separated on SDS-PAGE. Separated proteins were transferred to PVDF and probed with anti-HA to detects HA-PPK. Ponceau S is shown as a loading control. **B)** Ppx1 does not co-immunoprecipitate with Ddp1. 3HA-Ddp1 was immunoprecipitated using an anti-HA antibody and then recovered using protein A agarose beads before elution in sample buffer. Eluted proteins from indicated strains were separated on SDS-PAGE and transferred to PVDF membrane prior to probing with anti-HA to detect 3HA-Ddp1 and anti-Ppx1 to detect Ppx1. Images representative of n = 3 experiments for (A) and n = 2 for (B).

**Supplementary Figure 5.**
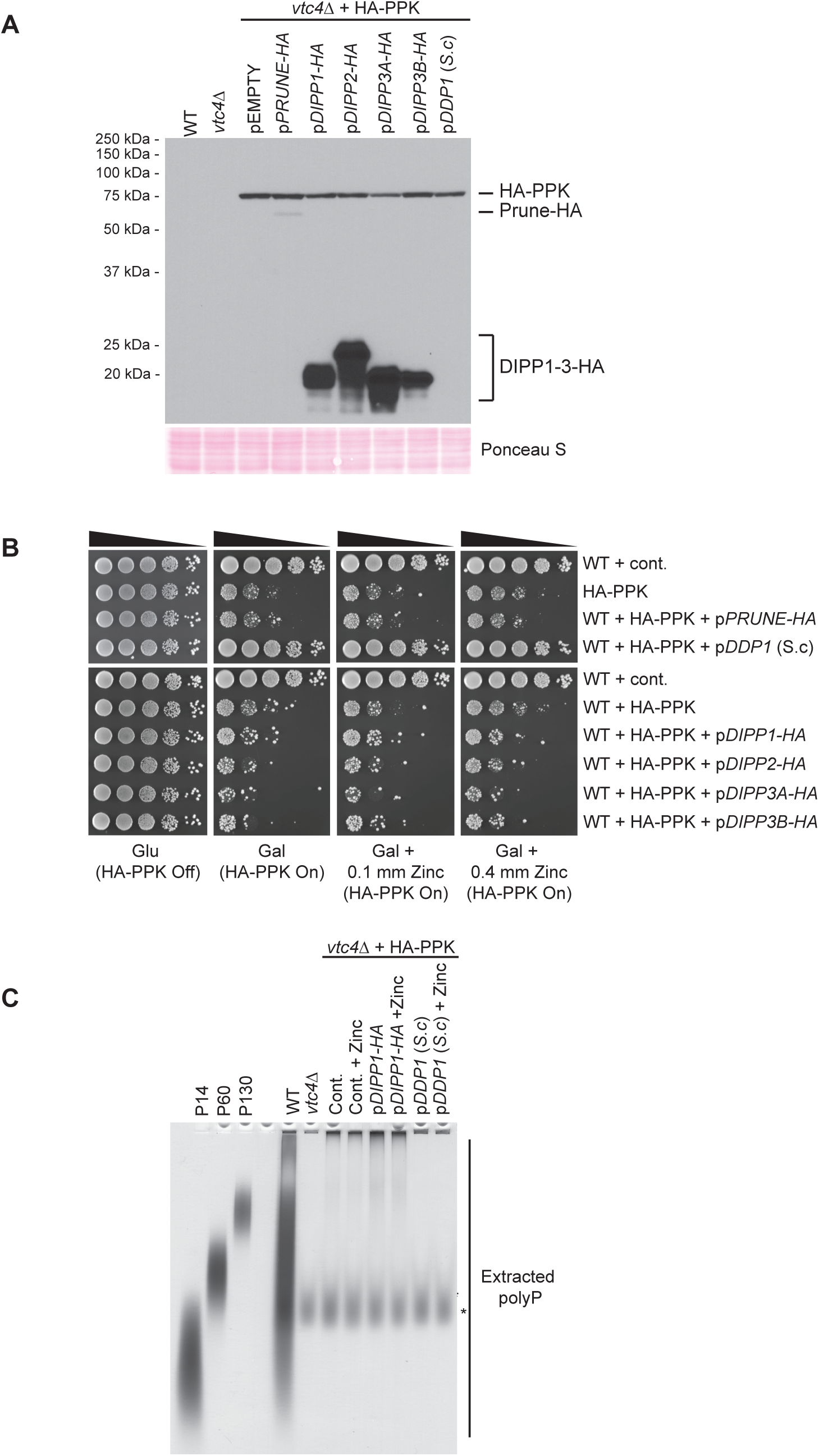
Overexpression of potential mammalian polyphosphatases. **A)** Protein extracts from the indicated strains were analyzed using a 12 % SDS-PAGE gel following a TCA-style lysis, prior to transfer to PVDF membrane and detection with anti-HA to detect HA-PPK. Strains are transformed with empty vector, HA-tagged PRUNE, HA-tagged DIPP1, HA-tagged DIPP2, HA-tagged DIPP3A, HA-tagged DIPP3B, or yeast Ddp1 expressing plasmids. Ponceau S stain is shown as a loading control. **B)** Strains with the indicated plasmids expressing potential mammalian polyphosphatases or yeast Ddp1 were spotted in 10-fold serial dilutions on indicated media supplemented with or without zinc sulfate. **C)** Polyphosphate was extracted from strains expressing the indicated plasmids and visualized using toluidine blue staining following separation on a 15.8 % acrylamide TBE-Urea gel. The asterisk indicates non-specific staining of an unknown product that is not polyP. Images representative of n = 3 experiments.

## SUPPLEMENTARY TABLES

**Supplementary Table S1 – Mass spectrometry data (.xlsx)**

**Supplementary Table S2 – Summary of MS dataset confirmations (.xlsx)**

**Supplementary Table S3 – Yeast strains used in this work (.xlsx)**

**Supplementary Table S4 – Antibodies used in this work (.xlsx)**

**Supplementary Table S5 – Plasmids used in this work (.xlsx)**

